# A fatal case of peritonitis caused by *Dysgonomonas capnocytophagoides* harboring the novel metallo-beta-lactamase gene *bla*_*DYB-1*_

**DOI:** 10.1101/2024.03.18.585585

**Authors:** Kazuo Imai, Masahiro Kodana, Ryuha Omachi, Tsutomu Inoue, Hirokazu Okada, Takuya Maeda

**Author notes:** **Corresponding author:** Kazuo Imai, MD, PhD, Department of Clinical Laboratory Medicine, Saitama Medical University, 38, Morohongo, Moroyama-machi, Iruma-gun, Saitama, Japan.

## Abstract

*Dysgonomonas capnocytophagoides* shows multidrug resistance to antibiotics and causes opportunistic infections in immunocompromised hosts. We report a fatal case of peritonitis caused by *D. capnocytophagoides* in Japan. We identified a novel chromosomally encoded class B1 metallo-beta-lactamase gene designated *bla*_*DYB-1*_ and an *ermFS* gene that contributed to multidrug resistance.

## Main text

*Dysgonomonas capnocytophagoides*, which belongs to a group of facultative anaerobic Gram-negative coccobacilli, is rarely isolated from stool and urine specimens of immunocompromised hosts (1, 2) and causes opportunistic infections (3-8). Antibiotic treatment is challenging because *D. capnocytophagoides* shows resistance to a broad range of beta-lactams, aminoglycosides, macrolides, and fluoroquinolones (4-8). However, the drug resistance mechanisms of *D. capnocytophagoides* have not yet been identified. Here, we present a fatal case of peritonitis due to multidrug-resistant *D. capnocytophagoides* that harbored a novel chromosomally encoded class B1 metallo-beta-lactamase (MBL) gene designated *bla*_*DYB-1*_ and an *ermFS* gene.

### The study

An 88-year-old Japanese woman with rheumatoid arthritis and chronic kidney disease was admitted to an intensive care unit because of colon perforation in 2023 in Japan. On the day of admission, she underwent surgery for colon perforation and was administered meropenem, vancomycin, and caspofungin. She developed a prolonged fever and required catecholamine blood pressure support. At 4 days after admission, metronidazole was started for secondary peritonitis and 2 sets of blood cultures were taken. Unfortunately, she died of septic shock and respiratory failure at 6 days after admission. Postmortem, the 2 sets of aerobic and anaerobic blood cultures were found to be positive for Gram-negative coccobacilli.

The isolate was grown on 5% sheep blood agar for 2 days in anaerobic conditions, and identified as *D. capnocytophagoides* by matrix-assisted laser desorption/ionization time-of-flight mass spectrometry (score, 2.080). Antibiotic susceptibility testing, conducted by the broth microdilution method using a Kyokuto Optopanel MP (Kyokuto Pharmaceutical, Tokyo, Japan) and MicroScan MF6J (Beckman Coulter, Brea, CA), found the isolate to be resistant to a broad range of beta-lactams and clarithromycin, ciprofloxacin, levofloxacin, and clindamycin (Table 1). The sodium mercaptoacetic acid test (9) showed the presence of an inhibition halo between the antibiotic discs (meropenem and ceftazidime) and the sodium mercaptoacetic acid discs (zone diameter > 15 mm), suggesting class B MBL production.

**Table 1.** Antimicrobial susceptibility testing of our isolate.

| Antibiotic | MIC<br>($\mu\text{g/ml}$) |
| --- | --- |
| Ampicillin | >2 |
| Sulbactam/ampicillin | >32 |
| Piperacillin | >128 |
| Tazobactam/piperacillin | >128 |
| Ceftriaxone | >64 |
| Cefotaxime | >64 |
| Cefmetazole | >64 |
| Imipenem/cilastatin | >16 |
| Meropenem | >16 |
| Ciprofloxacin | >1 |
| Levofloxacin | >2 |
| Moxifloxacin | 1 |
| Minocycline | 1 |
| Vancomycin | >4 |
| Metronidazole | $\leq 1$ |
| Clindamycin | >8 |
| Chloramphenicol | 4 |
| Sulfamethoxazole-trimethoprim | $\leq 0.5$ |

MIC, minimum inhibitory concentration.

We conducted whole genome sequencing by using the Flongle FLO-FLO114 (Oxford Nanopore Technologies, Oxford, UK) and iSeq100 instrument (Illumina, San Diego, CA). The sequence reads were assembled into 4,781,552 bp and 2,784 bp circular contigs, which were identified as a chromosomal genome and plasmid, respectively (GenBank accession number: AP028867 and AP028868). The 16S rRNA gene of the isolate had the closest relationship with that of *D. capnocytophagoides* type strain DSM22835 (GenBank accession number: NR_113133), with 97% coverage and 99.8% identity. Phylogenetic tree analysis based on 16S rRNA sequences also showed that our isolate belonged to *D. capnocytophagoides* (Supplementary Figure 1). DFAST and ResFinder indicated that our isolate harbored a chromosomally encoded putative MBL gene (designated *bla*_*DYB-1*_) and a chromosomally encoded *ermFS* gene (10). A consensus *Bacteroides* promoter sequence (11) was identified in the region immediately upstream of the putative MBL gene (Figure 1). A similar MBL gene (100% coverage and 98.8% identity) was also found in an assembled contig with a length of 172,569 bp in DSM22835 (GenBank accession number: AUFL00000000) and an assembled contig with a length of 457,687 bp in MRSN571793 (GenBank accession number: NZ_SOML00000000), and the genetic structure surrounding the MBL genes was highly conserved among our isolate, DSM22835, and MRSN571793 (Figure 1). No mobile element was identified around the MBL genes in TnCentral. The *ermFS* gene of the isolate was located on transposon Tn4551, which was identified in *Bacteroides fragilis* (GenBank: M17808).

**Figure 1.**
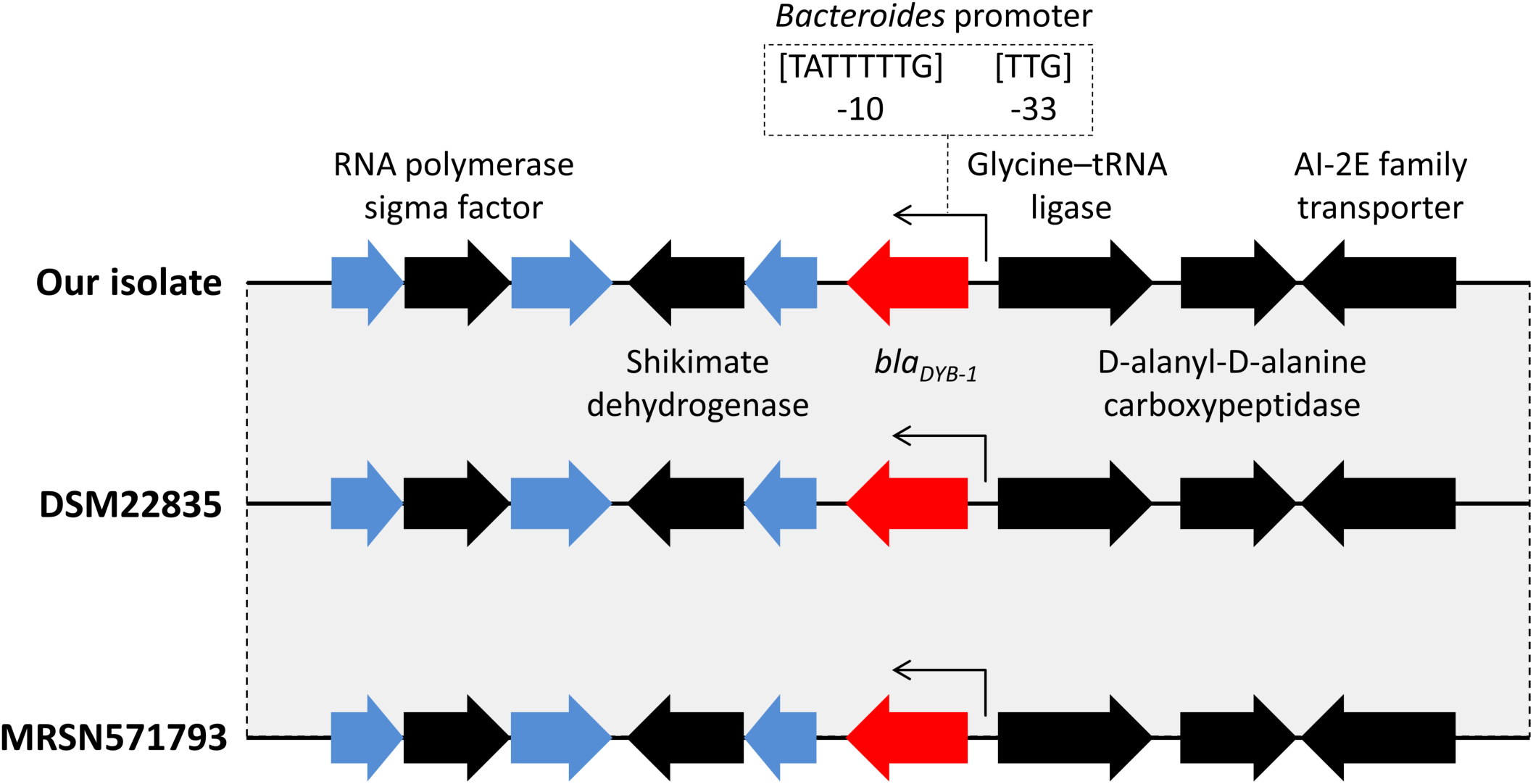
Genetic structure of isolates surrounding *bla*_*DYB-1*_. Blue arrows indicate putative proteins and gray shading shows >99% sequence identity among the isolates.

Based on amino acid sequence analysis, DYB-1 had the closest relationship with *B. fragilis* CfiA (12), a class B1 MBL, and DYB-1 harbored conserved zinc-interacting residues (His116, His118, Asp120, His196, Cys221, and His263) that are characteristic of class B1 MBLs (13, 14) (Supplementary Figure 2). SignalIP identify a typical N-terminal signal peptide (Supplementary Figure 2). The structure of the full-length MBL was predicted by AlphaFold2 on the Colab server, and was compared with the structure of CfiA (PDB accession number: 1ZNB). The conformation of the DYB-1 zinc-interacting active site was very similar to that of CfiA, suggesting that its enzymatic activity was di-zinc-dependent (Figure 2).

**Figure 2.**
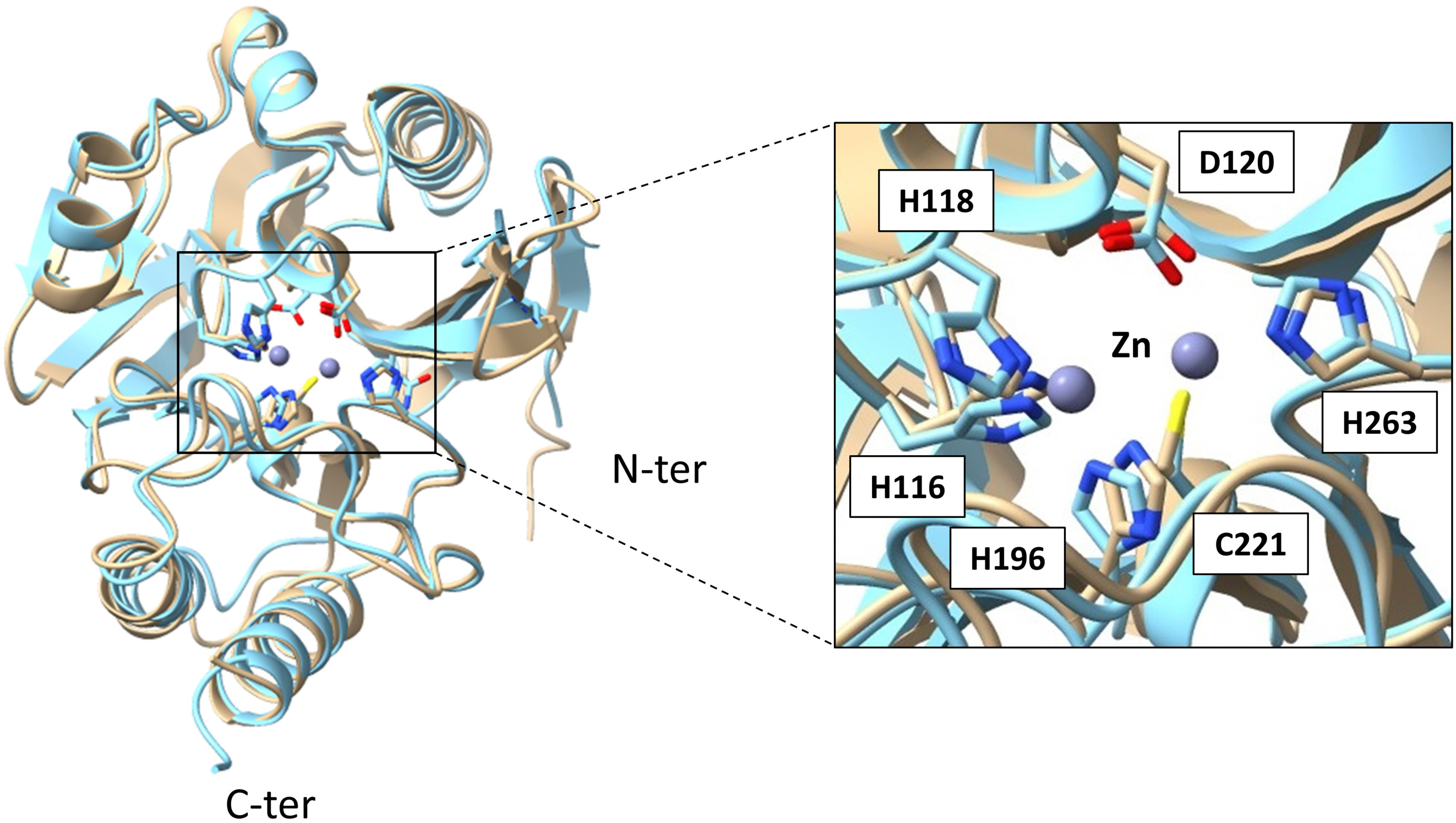
Structural analysis of DYB-1. Structural comparison of active sites between CfiA (blue) and DYB-1 (orange). The two zinc ions of the active site are indicated by gray circles.

Further, *bla*_*DYB-1*_ without signal peptide was amplified from our isolate by PCR and cloned into the pHSG398 vector (Takara Bio, Inc., Shiga, Japan) using the InFusion cloning method (Supplementary Table 1). Recombinant plasmids (pHSG398/*bla*_*DYB-1*_) and pHSG398 were transformed into the *Escherichia coli* TOP10 strain. Antibiotic susceptibility testing was conducted by the broth microdilution method using a MicroScan Neg 3J and Pos 2J (Beckman Coulter, Brea, CA). TOP10 cells transformed with pHSG398/*bla*_*DYB-1*_ showed a significant increase in the minimum inhibitory concentration (MIC) of beta-lactams, while there was no difference in the MIC of TOP10 cells transformed with pHSG398 only (Table 2).

**Table 2.** Antimicrobial susceptibility testing in transformation experiments.

| Antibiotic | MIC (µg/ml) |  |  |
| --- | --- | --- | --- |
|  | TOP10<br>pHSG398/*bla<sub>DYB-1</sub>* | TOP10/pHSG398 | TOP10 |
| Oxacillin | >4 | >4 | >4 |
| Ampicillin | >16 | ≤8 | ≤8 |
| Clavulanate/amoxicillin | ≤8 | ≤8 | ≤8 |
| Cefaclor | >16 | ≤8 | ≤8 |
| Cefazolin | 16 | ≤2 | ≤2 |
| Cefoxitin | ≤4 | ≤4 | ≤4 |
| Cefotaxime | 32 | ≤1 | ≤1 |
| Ceftazidime | >16 | ≤1 | ≤1 |
| Cefepime | ≤2 | ≤2 | ≤2 |
| Imipenem/cilastatin | 2 | ≤1 | ≤1 |
| Meropenem | 8 | ≤0.12 | ≤0.12 |
| Doripenem | >4 | ≤1 | ≤1 |
| Aztreonam | ≤4 | ≤4 | ≤4 |

MIC, minimum inhibitory concentration.

In this study, we identified the novel chromosomal class B1 MBL gene *bla*_*DYB-1*_ that contributes to resistance to a wide range of beta-lactams and an *ermFS* gene that contributes to resistance to macrolides and clindamycin in *D. capnocytophagoides*. Because there were no mobile elements around *bla*_*DYB-1*_, its risk of transmission between organisms may be low (15). However, given that foreign mobile elements may insert themselves at genomic locations around *bla*_*DYB-1*_, the transmission risk of MBL genes in hospital settings and immunocompromised hosts cannot be denied. In addition, *D. capnocytophagoides* could function as a reservoir for the *ermFS* gene through transposon Tn4551 among pathogens.

Prior case reports demonstrated that *D. capnocytophagoides* shows susceptibility to clindamycin, tetracycline, chloramphenicol, and sulfamethoxazole-trimethoprim (4-8). Our isolate showed susceptibility to moxifloxacin and metronidazole as well as tetracycline, chloramphenicol, and sulfamethoxazole-trimethoprim *in vitro*. The antibiotic resistance genes for those antibiotics were not found in our isolate. Taking genetic characteristics into account, those antibiotics are possible treatment options for *D. capnocytophagoides*. In our case, the early administration of metronidazole may have controlled the peritonitis caused by *D. capnocytophagoides*. Timely species identification and appropriate antibiotic treatment are crucial for the effective management of *D. capnocytophagoides* infection in immunocompromised hosts.

## Conclusions

We reported a fatal opportunistic infection caused by *D. capnocytophagoides* with the novel MBL gene *bla*_*DYB-1*_ and *ermFS* contributing to its multidrug resistance mechanisms. This study indicates the need for further surveillance of *D. capnocytophagoides* harboring *bla*_*DYB-1*_ focusing on its pathogenicity, clonal diversity, and role as an MBL gene reservoir.

## Acknowledgements

Not applicable.

## Conflict of Interest Statement

The authors declare that they have no conflicts of interests

## Funding

This work was partially supported by JSPS KAKENHI Grant Number JP21K07375.

## Data Availability

The assembled chromosomal genome and plasmid were deposited in GenBank (Accession number: AP028867 and AP028868).

## Supplementary Figure Legends

**Supplementary Figure 1**. Phylogenetic analysis based on the 16S rRNA gene sequence. The phylogenetic tree was constructed by the neighbor-joining method. The bootstrap tests (1,000 replicates) are shown next to the branches.

**Supplementary Figure 2**. Amino acid alignment of DYB-1 with other function-characterized class B beta-lactamases. The amino acid numbering was according to the scheme for class B beta-lactamases (Galleni M et al., *Antimicrob Agents Chemother*. 2001; 45(3): 660–663).

**Supplementary Figure 3**. Phylogenetic analysis of DYB-1 with other function-characterized class B beta-lactamases. The phylogenetic tree was constructed by the neighbor-joining method.

**Supplementary Table 1.**
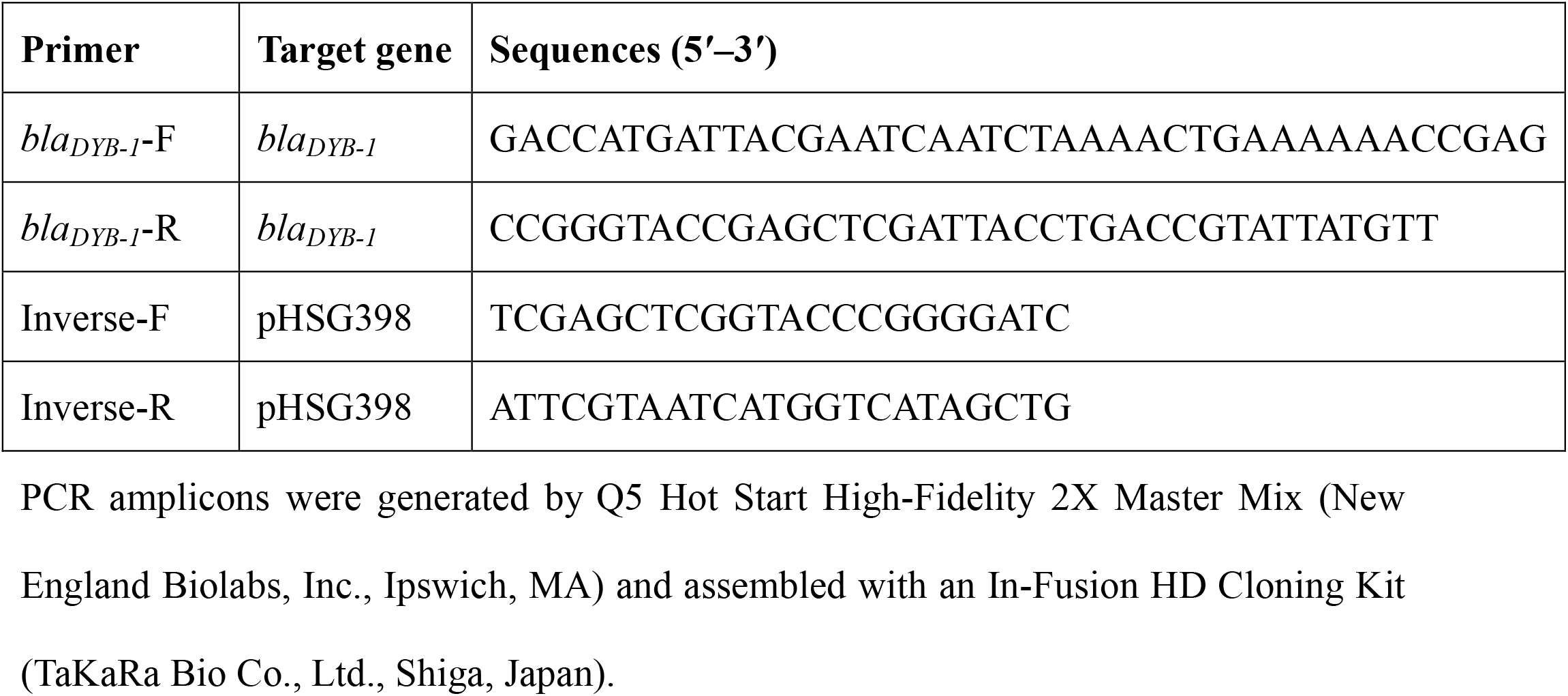
Primers for cloning the *bla*_*DYB-1*_ gene into pHSG398.

